# Adaptive optics two-photon endomicroscopy enables deep brain imaging at synaptic resolution over large volumes

**DOI:** 10.1101/2020.05.29.124727

**Authors:** Zhongya Qin, Congping Chen, Sicong He, Ye Wang, Kam Fai Tam, Nancy Y. Ip, Jianan Y. Qu

## Abstract

Optical deep brain imaging in vivo at high resolution has remained a great challenge over the decades. Two-photon endomicroscopy provides a minimally invasive approach to image buried brain structures, once it is integrated with a gradient refractive index (GRIN) lens embedded in the brain. However, its imaging resolution and field of view are compromised by the intrinsic aberrations of the GRIN lens. Here, we develop a two-photon endomicroscopy by adding adaptive optics based on the direct wavefront sensing, which enables recovery of diffraction-limited resolution in deep brain imaging. A new precompensation strategy plays a critical role to correct aberrations over large volumes and achieve rapid random-access multiplane imaging. We investigate the neuronal plasticity in the hippocampus, a critical deep brain structure, and reveal the relationship between the somatic and dendritic activity of pyramidal neurons.

## 1. INTRODUCTION

Advances in two-photon microscopy have greatly propelled the studies of neural circuits and brain functions in the past decades, by enabling high-resolution morphological and functional imaging in the living brain. In conjunction with various fluorescent proteins and indicators, two-photon microscopy allows for direct visualization of fine neuronal structures and continuous monitoring of dynamic neural activities at the spatiotemporal scale across orders of magnitude^1,2^. However, imaging has been restricted to the superficial brain regions because of the severe attenuation of both excitation and emission photons at depth caused by tissue scattering. Although a longer excitation wavelength or red-shifted fluorescence labeling can alleviate the scattering effect, the imaging depth is still limited to one to two millimeters in the mouse brain and the imaging quality degrades rapidly with increasing depth^3,4^.

To image deeper subcortical structures beyond this limit, an endomicroscopy has been adopted that relies on implanting a miniature gradient refractive index (GRIN) lens in the brain^5–7^. The rod-like GRIN lens acts as a relay between the microscope objective and the sample below. This lens has been integrated with the single-photon epifluorescence microscope to image various neurons lying deep in the brain that are beyond the reach of conventional microscopy^8–10^. However, the optical resolution for such a miniature single-photon microscope has been limited to the cellular level, with the image contrast reduced by the out-of-focus fluorescence background. Two-photon endomicroscopy incorporating a high numerical aperture (NA) GRIN lens has enabled resolving subcellular structures at high resolution^5,11,12^. However, the imaging field of view (FOV) is restricted to tens of microns in diameter due to the severe off-axis aberrations of the GRIN lens. Moreover, for three-dimensional (3D) imaging, because the GRIN lens is embedded in the biological sample, the focal plane is tuned by changing the distance between the microscope objective and the GRIN lens, which could lead to severe on-axis aberrations when the excitation focus deviates axially from the designed optimum^5^. Therefore, the enlarged and distorted point spread function (PSF) resulting from the intrinsic aberrations significantly limits the 3D imaging volume of GRIN lens-based endomicroscopy.

In recent years, adaptive optics (AO) has greatly advanced two-photon imaging by introducing a compensatory wavefront distortion to the excitation laser that cancels the system- or specimen-induced aberrations^13,14^. The wavefront distortion can be determined by either direct^15–19^ or indirect^20–22^ wavefront sensing. In previous studies, a sensorless AO approach based on pupil segmentation was used to correct the aberrations of the GRIN lens, improving the resolution over an enlarged imaging FOV of 205×205 μm^2^ in a fixed brain slice^23,24^. However, this method is time-consuming for wavefront estimation and sensitive to sample motion, which limits its application for *in vivo* imaging. An unmet challenge is to develop robust AO two-photon endomicroscopy enabling high-resolution imaging over large volumes in the living mouse brain.

Here we developed an AO two-photon endomicroscope based on direct wavefront sensing for high-resolution deep brain imaging *in vivo* (**Supplementary Fig. 1**). Our approach is to use the two-photon excited fluorescence (TPEF) signal as the intrinsic guide star inside biological tissues. The wavefront of the descanned guide star is averaged over a small region (typically 30×30 μm^2^) and measured with the Shack-Hartmann wavefront sensor (SHWS) consisting of a microlens array and an electron-multiplying charge-coupled device (EMCCD)^25^. In order to achieve fast AO correction over large volume, we first characterized the aberrations of the GRIN lens and developed a lookup table method to precompensate for its intrinsic aberrations before *in vivo* imaging. Direct wavefront sensing combined with lookup table method permits accurate, rapid and robust estimation of the optical aberrations during *in vivo* brain imaging^17^. The wavefront information is then used by the deformable mirror (DM) that forms a closed loop with the SHWS to create a compensatory distortion to the excitation light and yield a diffraction-limited focus inside the brain tissues. Using the lookup table and following *in situ* AO correction, we achieved structural imaging of hippocampal CA1 neurons at synaptic resolution over a large volume. We studied neuronal plasticity of the hippocampus under various pathological conditions. Furthermore, by combining our new endomicroscope with a fast electrically tunable lens, we demonstrated quasi-simultaneous multiplane calcium imaging of neuronal somata and dendrites at high spatiotemporal resolution.

## 2. RESULTS

To evaluate the efficacy of our approach, we first characterized the intrinsic aberrations of the GRIN lens using an *in vitro* setup. The GRIN lens was embedded in a rhodamine/agarose mixture that contains sparsely distributed and immobilized fluorescent beads (200-nm diameter). The aberration of the GRIN lens was measured by the wavefront distortion of rhodamine fluorescence, whereas the PSF was determined by imaging the individual fluorescent bead. To achieve the optimal imaging performance, the GRIN lens was precisely aligned to share the same optical axis with the objective. Considering the cylindrical symmetry of the GRIN lens, the cylindrical coordinates (r, θ, z) were adopted to describe the imaging location in the sample where z=0 corresponds to the designed focal plane of the GRIN lens. The representative on-axis and off-axis aberrations of the GRIN lens at various depths are shown in **Supplementary Figs. 2 and 3**, which are primarily dominated by spherical aberration and astigmatism respectively, consistent with a previous study^24^. As can be seen from the figures, despite the aberrations increasing rapidly and the PSF becoming severely distorted when the imaging position deviates from the center point, AO correction can effectively increase the fluorescence signal and recover the diffraction-limited performance at various depths (**Supplementary Figs. 2 and 3**).

We next applied the AO approach to the *in vivo* imaging of the mouse hippocampus, in which a subset of pyramidal neurons express green fluorescent proteins under thy1 promoter (Thy1-GFP mice). The CA1 pyramidal neurons are vertically oriented with a typical laminar organization spanning hundreds of microns in depth^26,27^. As can be seen from **Fig. 1a** and **Supplementary Fig. 3**, with system correction alone, the neuronal somata and dendrites are severely blurred. After full AO correction and subsequent deconvolution from the local PSF, however, not only is the fluorescence intensity drastically improved, the dendrites and spines can also be clearly visualized owing to the substantially enhanced resolutions (**Fig. 1b** and **Supplementary Fig. 4**). The corrective wavefront primarily contains astigmatism as previously characterized for the off-axis scenario, implying the aberrations mainly came from the imperfections of the GRIN lens (**Fig. 1c**). The spectral power shows that the higher spatial frequency can be significantly recovered by applying full AO correction with subsequent deconvolution (**Fig. 1d**). Quantitative measurement from individual dendritic spines shows that both lateral and axial resolutions approached diffraction-limited performance for the GRIN lens with NA of 0.8 (**Figs. 1e-f**). Furthermore, even for the on-axis scenario where the GRIN lens was designed to yield the best performance, we still achieved significant improvement in resolution after full AO correction, with dendritic spines as deep as 300 μm clearly resolved (**Supplementary Fig. 5**). These results demonstrate that AO based on direct wavefront sensing can effectively correct the aberrations of the GRIN lens and recover the diffraction-limited resolution during *in vivo* imaging.

**Figure 1.**
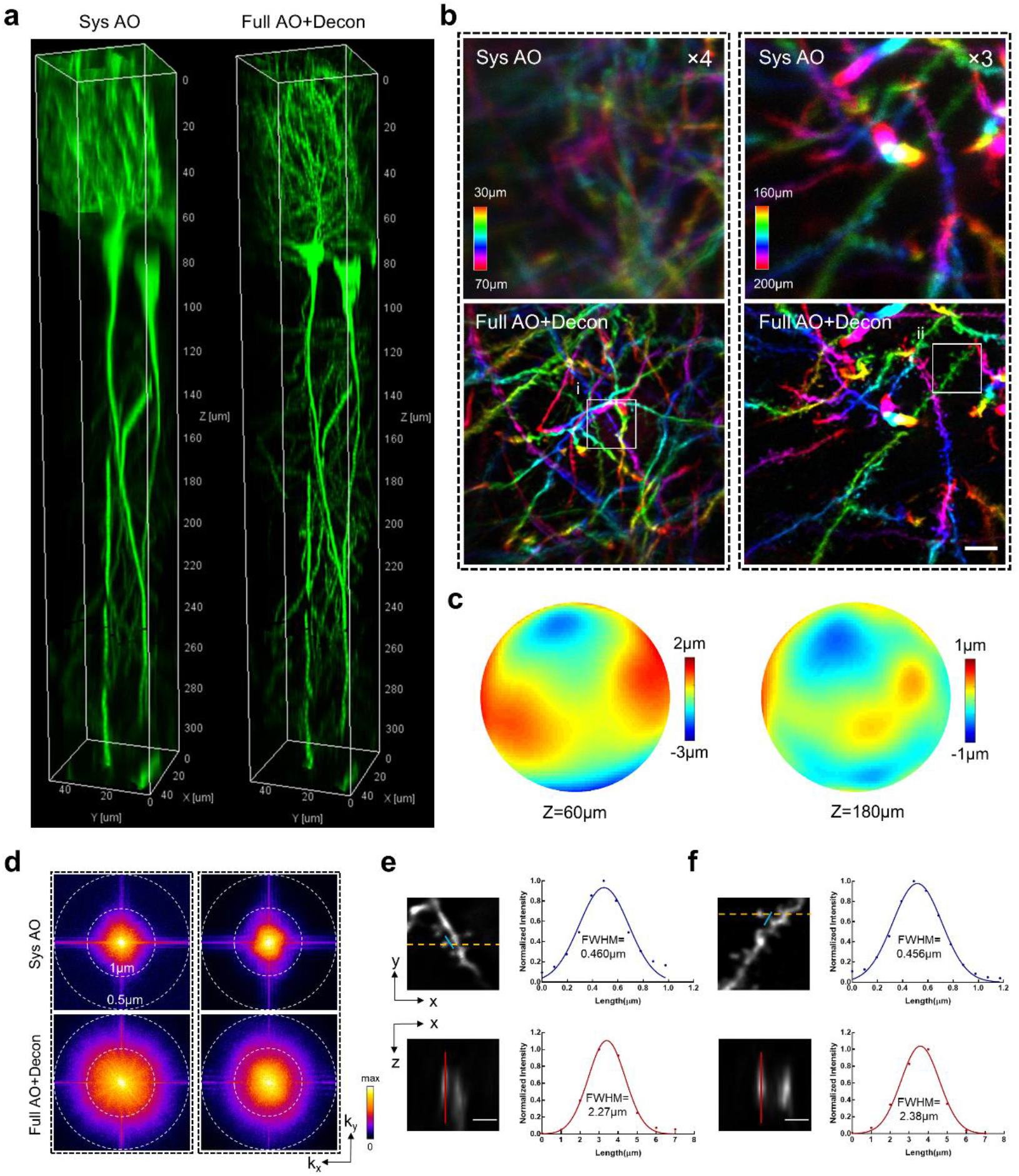
Direct-wavefront-sensing AO effectively restores diffraction-limited resolution at depth during *in vivo* brain imaging. (a) 3D reconstruction of a column (center located at r=60 μm) of hippocampal CA1 pyramidal neurons in Thy1-GFP mice imaged with our two-photon endomicroscope with system correction only (left) and with full correction plus subsequent deconvolution (right). Full AO correction is performed every 30 μm of depth. (b) Depth-color-coded xy maximum-intensity projection (MIP) of the stack images (left column: 30-70μm, right column: 160-200 μm) from 3D images in Fig. 1(a). The images with system correction have been digitally enhanced fourfold and threefold as indicated for better visualization. Scale bar: 5 μm. (c) Corrective wavefronts of the DM used for full AO correction of the stack images in (b). (d) Spectral power in spatial frequency space (kx, ky) for the images in (b). (e-f) Magnified views of the dendritic spines corresponding to the boxed regions (i-ii) in (b). The spines are shown in lateral (xy) and axial (xz) view. The axial view is shown through the plane defined by the yellow dashed line. Intensity profiles along the blue and red lines are plotted with the curve fitted by a Gaussian function. Scale bar: 2 μm.

Next we sought to investigate the extent to which our AO approach can enlarge the imaging FOV *in vivo*. Because the aberrations vary from site to site, we performed AO correction in a number of subregions of 50×50 μm^2^ each separated by a distance of 30 μm laterally and stitched these subregions together to form a high-resolution image of large FOV. Considering that the photon budget and fluorescence may be too weak for wavefront measurement, we propose a lookup table method to precompensate for the aberrations in wavefront measurement (**Supplementary Fig. 6**). We found that the corrective wavefront from the lookup table serves as a close approximation of the optimal wavefront measured *in vivo* (**Fig. 2**). The lookup table method can greatly boost the fluorescence signal above and beyond system correction and thus significantly decrease the exposure time required for direct wavefront measurement. Due to the large off-axis aberrations of the GRIN lens, the imaging FOV with only system correction was restricted to a small central region (<100 μm in diameter) of poor resolution (**Fig. 3a** and **Supplementary Fig. 7a**). After AO full correction, the effective FOV was enlarged to 300 μm together with the recovery of synaptic resolution. The neuronal structures including somata, dendrites and individual spines can be clearly visualized across the entire FOV, covering all layers of the hippocampus CA1 region (**Fig. 3b** and **Supplementary Fig. 7b**).

**Figure 2.**
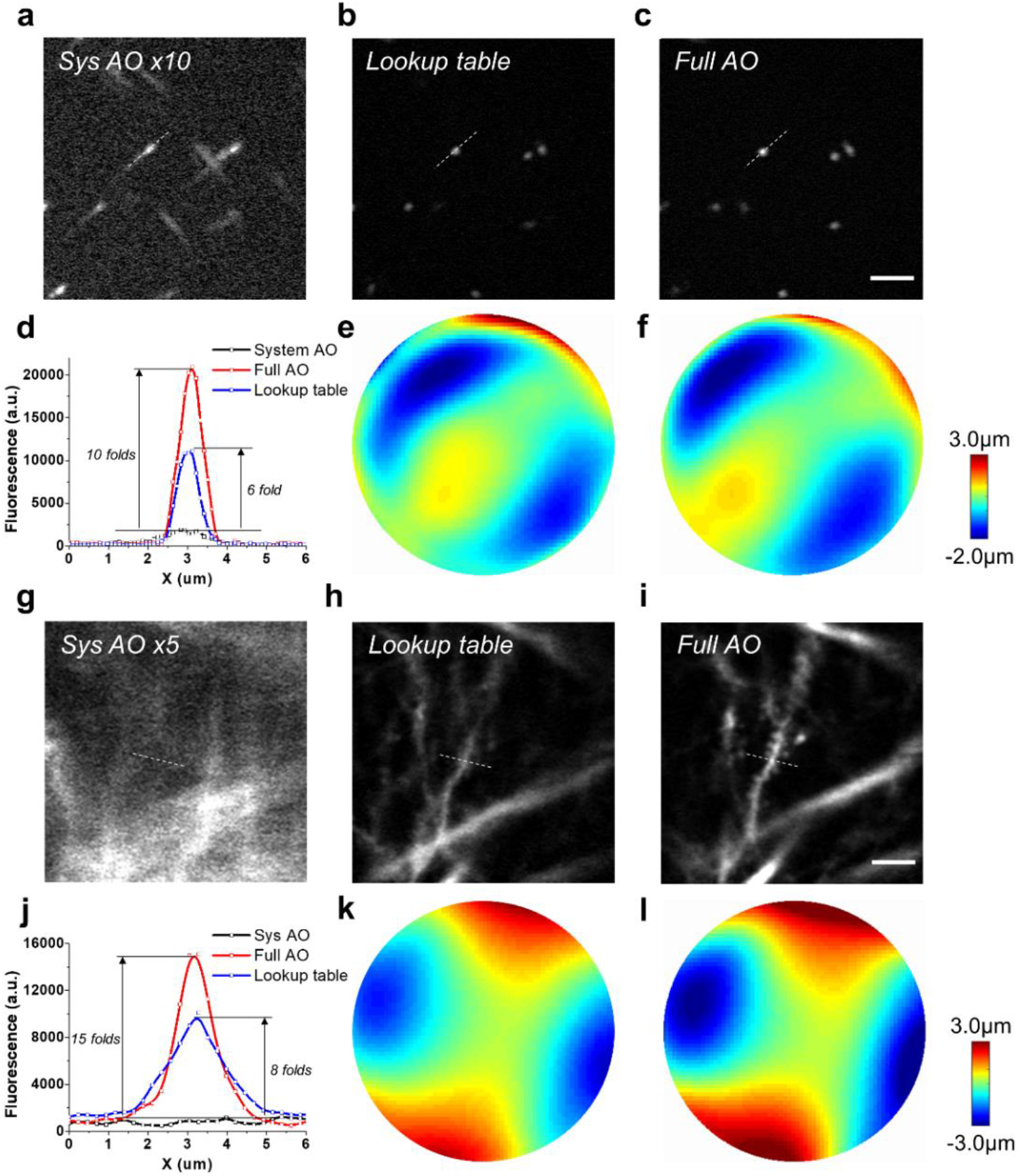
Lookup-table-based precompensation for GRIN lens aberrations. **(a-f)** *In vitro* imaging using fluorescent beads of 0.2 μm in diameter. **(a-c)** Fluorescent images with system AO correction (a), lookup table correction (b) and full AO correction (c). **(d)** Intensity profile along the dashed lines in (a-c). **(e-f)** The corrective wavefront used in lookup table correction (b) and full AO correction (c). **(g-l)** *In vivo* imaging of hippocampal neurons in mice. **(g-i)** Fluorescent images with system AO correction (g), lookup table correction (h) and full AO correction (i). Scale bar: 5 μm. **(j)** Intensity profile along the dashed lines in (g-i). **(k-l)** The corrective wavefront used in lookup table correction (h) and full AO correction (i).

**Figure 3.**
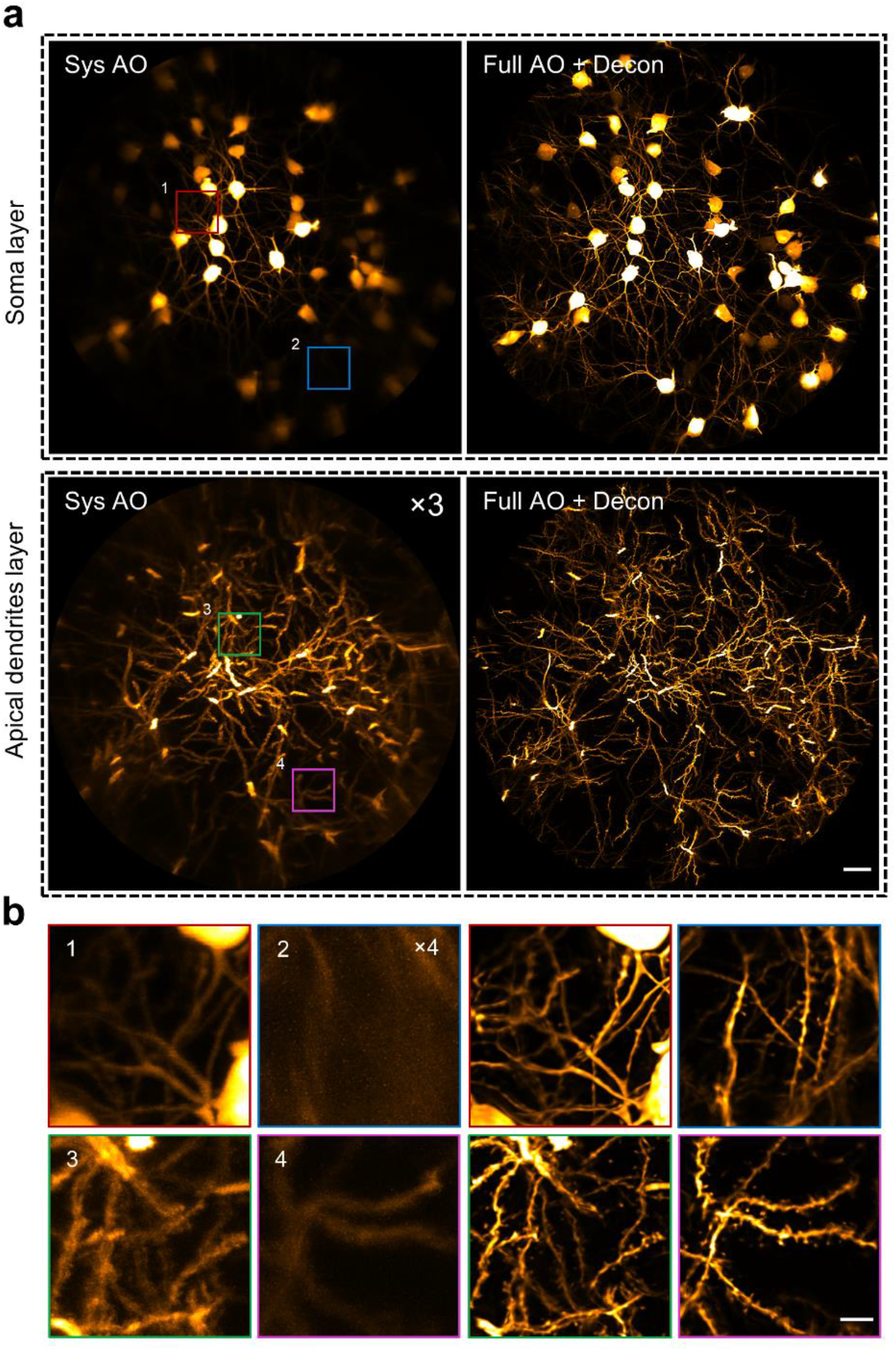
AO two-photon endomicroscope enables *in vivo* imaging of the mouse hippocampus at synaptic resolution over a large FOV. (a) MIP images of different layers of hippocampal CA1 pyramidal neurons in Thy1-GFP mice with only system correction (left column) and with full correction plus subsequent deconvolution (right column). Depth range of projection for soma layer: 90-120 μm; apical dendrite layer: 160-190 μm. The images with system correction were enhanced for better visualization. Scale bar: 20 μm. (b) Left: four magnified views of the sub-regions indicated by the numbered boxes in (a). Right: views of corresponding sub-regions with full AO corrections. Scale bar: 5 μm.

Taking advantage of our approach, we further investigated the neuronal plasticity of the hippocampus under various pathological conditions via *in vivo* time-lapse imaging of the dendritic and spine dynamics. We first demonstrated the dendritic alteration in response to the neuronal injury induced by a laser-mediated microsurgery. We found that micro-lesion of single dendritic shaft will trigger recurrent spinogenesis in the neighboring regions, whereas cutting the dendritic branches with laser will lead to prolonged degeneration of the injured dendrites (**Supplementary Fig. 8**). In addition, we also monitored the spine dynamics in mice with kainic acid-induced epileptic seizure. In contrast to the dynamic neuronal changes induced by laser microsurgery, our findings show that the changes in spine stability caused by kainic acid-induced epileptic seizure are insignificant, which is in agreement with another study of the hippocampus of mice experiencing pilocarpine-induced epileptic seizure^28^. These results shed light on the neuronal mechanisms underlying neurodegenerative disorders.

In addition to morphological imaging, we further applied the AO two-photon endomicroscope to the *in vivo* functional calcium imaging of pyramidal neurons in the hippocampus of awake behaving mice that expressed a calcium indicator (AAV-CaMKII-GCaMP6s). Taking advantage of the large imaging volume provided by AO correction, we developed a random-access multiplane imaging method that can quickly capture arbitrarily selected regions of interest (ROIs) across the effective imaging space (**Fig. 4a**). The galvanometer scanners were programmed to scan the ROIs sequentially and an electrically tunable lens (ETL) was synchronized with the scanners for fast switching between imaging planes (**Fig. S9**). To recover the optimal imaging performance in all the ROIs, the wavefront distortion in each ROI was measured individually and the corrective pattern of DM was synchronously updated to compensate for the field-dependent aberrations (**Supplementary Fig. 9**). After AO full corrections in ROIs, the imaging system can clearly resolve the tightly-packed neurons with the calcium transients accurately recorded and without interference from the neighboring neurons (**Fig. 4d-f**) Moreover, the fine neurites and individual spines contaminated by the neuropil background can be easily distinguished owing to the greatly enhanced resolution and fluorescence intensity (**Figs. 4b-c**). The results demonstrate that our AO approach provides an accurate and sensitive characterization of neuronal activity and enables simultaneous calcium imaging of neuronal somata and dendrites at synaptic resolution.

**Figure 4.**
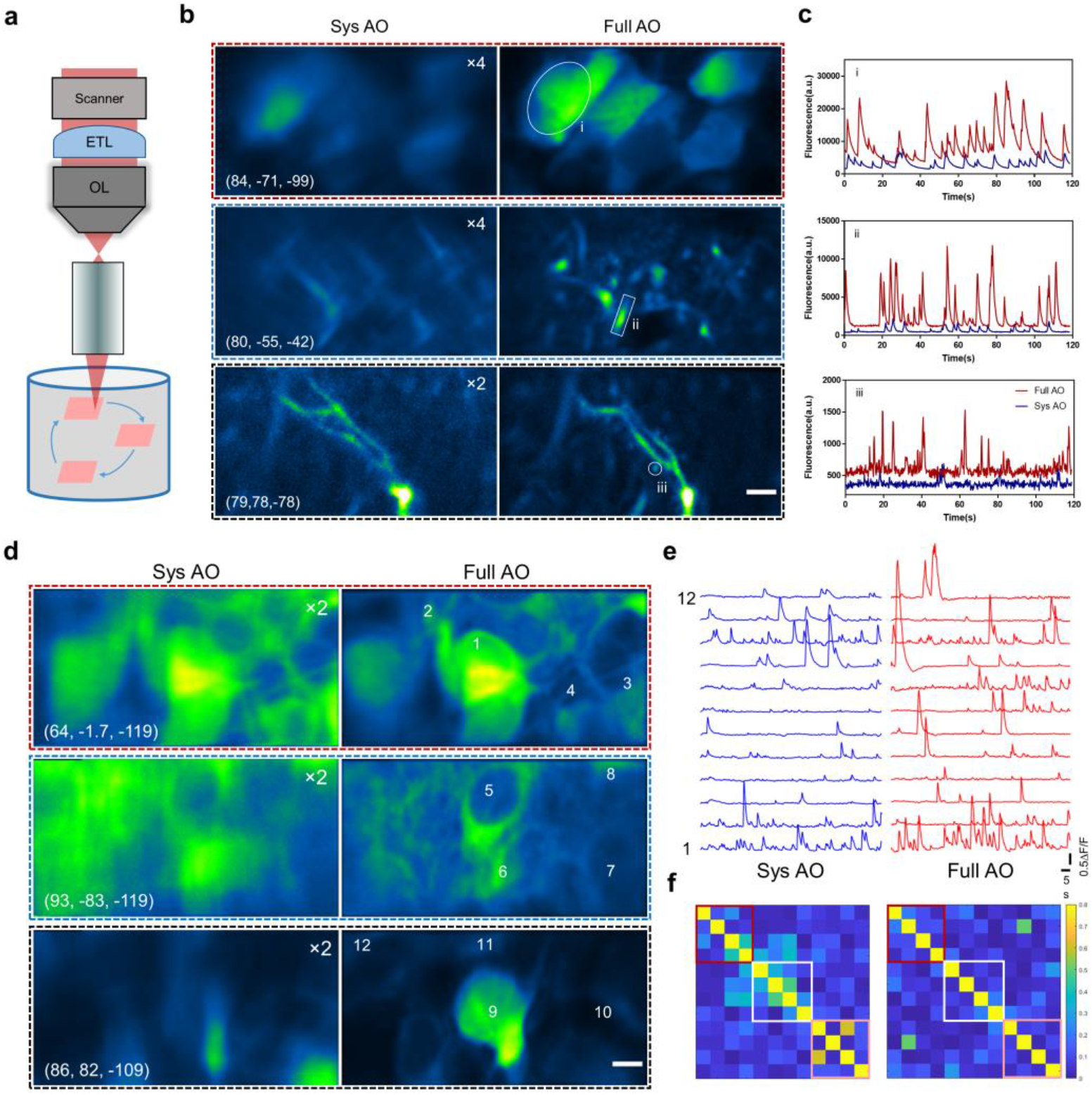
Random-access multiplane Ca^2+^ imaging of hippocampal neurons *in vivo*. (a) Schematic diagram of random-access multiplane imaging. Three image planes over the entire imaging volume of 0.3×0.3×0.3 mm^3^ can be randomly selected and sequentially scanned at 5 Hz with synchronized ETL and xy galvanometer scanners. (b) Multiplane Ca^2+^ imaging of neuronal somata, dendrites and spines at various locations with system correction (left) and with full correction (right). The virus AAV-CaMKII-GCaMP6s was injected into the hippocampus CA1 of C57/B6 mice. Images are shown as average-intensity projection of 600 frames. The images with system correction were enhanced as indicated for better visualization. The cylinder coordinates: (μm, deg, μm). Scale bar: 5 μm. (c) Fluorescence traces for the ROIs indicated in (b). (d) Ca^2+^ imaging of pyramidal neurons in hippocampus CA1 with system and full AO correction. Images are shown as average-intensity projections of 600 frames. Scale bar: 5μm. (e) Calcium transients (ΔF/F) of the selected neurons as shown in (a). (f) Correlation coefficient matrices calculated from ΔF/F traces of all neurons in (b). Colored boxes refer to the three ROIs as shown in (a).

We then applied this AO-assisted functional imaging technology to investigate the relationship between somatic and dendritic activity in mouse hippocampus CA1. The soma and two apical dendrites of single pyramidal neuron were targeted for simultaneous calcium imaging (**Figs. 5a-b**). We found that the dendritic and somatic signals are highly correlated, despite their amplitudes and kinetics are largely different (**Fig. 5c-e**). This functional correlation of dendrites and neuronal somata has also been reported in the cortex of awake mouse, in which the persistent coupling of somato-dendritic activity is unchanged by stimuli and mouse locomotion^29^. Our further results show that the correlated somato-dendritic activity in mouse hippocampus is brain-state dependent. In compared with full wakefulness, isoflurane-induced light anesthesia will not only weaken the activity of different neuronal compartments but also disrupt the somato-dendritic correlation (**Supplementary Fig. 10**). In future work, it would be of great interest to extend this method to study the dendritic integration in awake mice performing behavioral tasks^30,31^.

**Figure 5.**
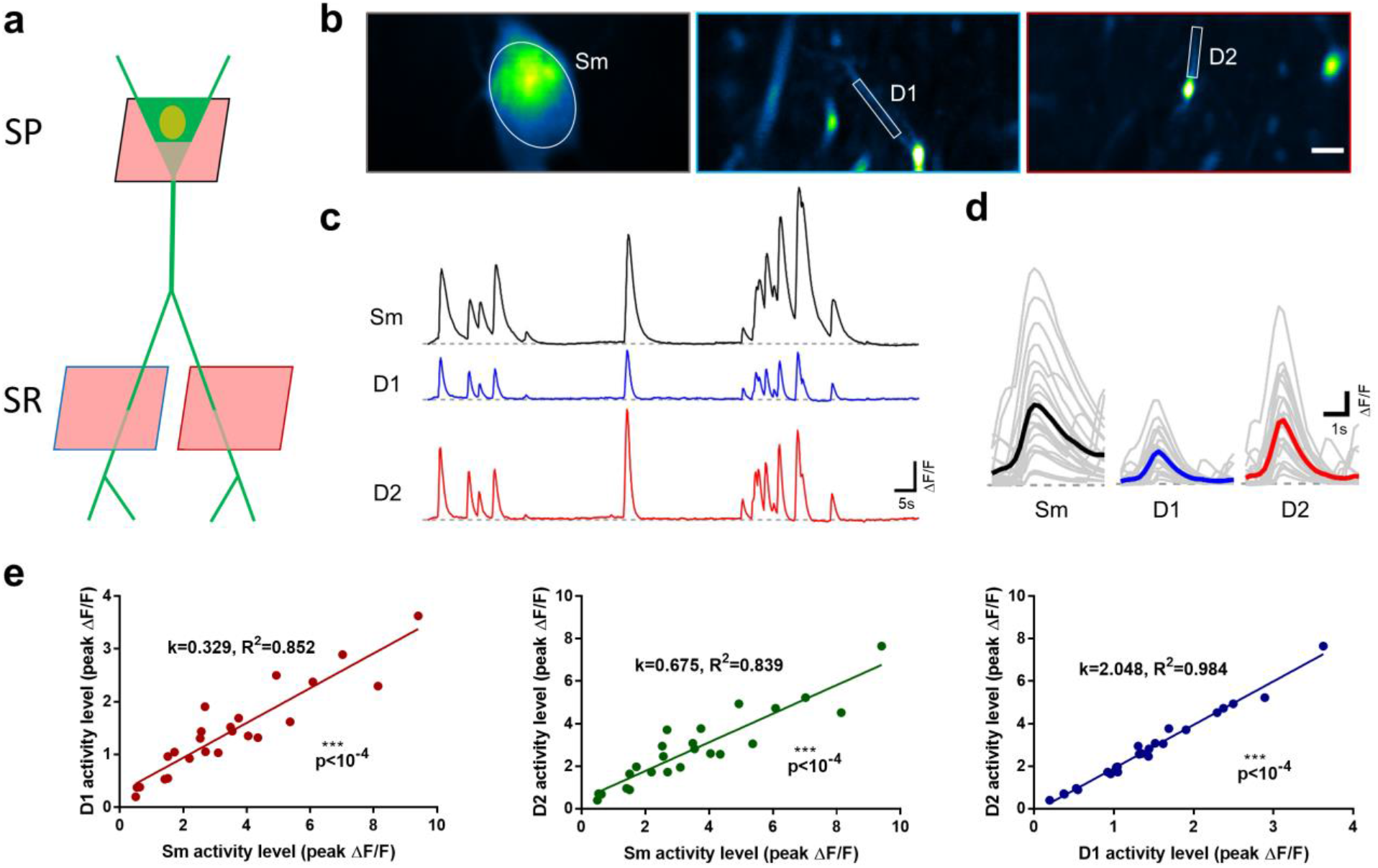
AO-assisted multiplane Ca^2+^ imaging of somato-dendritic activity in hippocampus CA1. (a) Experimental approach to target the dendrites and soma for single neuron recording in hippocampus CA1. SP: stratum pyramidale, SR: stratum radiatum. (b) Quasi-simultaneous Ca^2+^ imaging of spontaneous activity from the soma (Sm) and two dendrites (D1, D2) in awake behaving mice. Images are shown as standard-deviation (STD) projection of 600 frames. Scale bar: 5μm. (c) Calcium transients (ΔF/F) of the soma and dendrites as shown in (b). (d) Firing events of soma (Sm) and dendrites (D1, D2) as shown in (b). Grey and colored curves represents the individual and average event, respectively. (e) Relationship between activity strength of the soma-dendrite pairs and dendrite-dendrite pair.

## 3. CONCLUSION

In this work, we combined the AO technique based on the direct wavefront sensing of the TPEF guide star with a high-NA endomicroscope, which enables the recovery of diffraction-limited resolution over large imaging volumes. Using this system, we achieved *in vivo* morphological imaging of pyramidal neurons at synaptic resolution across all layers of mouse hippocampus CA1. Moreover, by integrating the system with a random-access multiplane imaging technique, we demonstrated quasi-simultaneous calcium imaging of separately distributed somata and the dendrites in the hippocampus of awake behaving mice. The AO endomicroscope can also benefit the imaging of other deep brain structures such as the striatum, the substantia nigra and the hypothalamus^32^. It should be noted that although brain tissue can be surgically removed and replaced with a cannula window^33^ or glass plug^34^ to provide direct optical access, the procedure requires removing too much tissue and is not applicable to deep brain imaging. An endomicroscope based on a miniature GRIN lens provides an approach to deep brain imaging with minimized trauma, but suffers from poor resolutions due to the inherent aberrations of the GRIN lens. One way to solve this problem directly is by adding an extra lens inside the GRIN lens assembly to correct the off-axis aberrations^35^. However, the GRIN lens is only optimized at the designed working distance. For *in vivo* imaging where the GRIN lens remains fixed in the biological samples, 3D imaging is achieved by changing the laser convergence entering the GRIN lens, which could lead to severe aberrations when the focus deviates from the designed working plane. Our AO endomicroscope, on the other hand, provides a versatile approach for correcting the aberrations of various GRIN lenses with suboptimal design. This study demonstrates the great potential of the AO two-photon endomicroscope to facilitate neuroscience research in the deeper regions of the brain.

## Acknowledgements

This work was supported by the Hong Kong Research Grants Council through grants 662513, 16103215, 16148816, 16102518, T13-607/12R, T13-706/11-1, T13-605/18W, C6002-17GF, C6001-19EF, N_HKUST603/19 and the Innovation and Technology Commission (ITCPD/17-9), and the Area of Excellence Scheme of the University Grants Committee (AoE/M-604/16, AOE/M-09/12) and the Hong Kong University of Science & Technology (HKUST) through grant RPC10EG33

## Author contributions

Z.Q., C.C. and J.Y.Q. conceived of the research idea. C.C. and Z.Q. designed and conducted the experiments and data analysis. Z.Q., S.H. and C.C. built the AO two-photon and multiplane imaging system. Y.W. and K.F.T. carried out the surgery of virus injection and GRIN lens implantation. C.C. and Z.Q. took the lead in writing the manuscript with inputs from all other authors.

## Methods

### Adaptive optics two-photon endomicroscopy

The schematic of our home-built AO two-photon endomicroscopy system is shown in **Supplementary Fig. 1**. The beam of a tunable mode-locked femtosecond laser (Coherent, Mira 900) was expanded by a pair of achromatic lens to slightly overfill the aperture of the DM (Alpao, DM97-15). The DM was conjugated to the 5 mm galvanometric *x*-scanning mirror (Cambridge Technology, 6215H) by a 4*f* telescope formed by two VIS-NIR achromatic doublets L3 and L4 (Edmunds, 49-365 and 49-794). Two scanning mirrors were mutually conjugated by the lens pair L5 and L6, each of which consisted of two doublets (Edmunds, 49-392). Finally, the galvanometric *y*-scanning mirror was conjugated to the back focal plane of the objective by a 4*f* relay formed by the scan lens L7 (consisting of two doublets (Edmunds, 49-391)) and the tube lens L8 (Edmunds, 49-393). A 10× air objective (Carl Zeiss, Plan-Apochromat, NA=0.45) was used to match the image NA of the GRIN lens (GRINTECH GmbH, GT-MO-080-0415-810, image NA=0.415, object NA=0.8) and mounted on an encoded translation stage (Thorlabs, LNR50SEK1) for axial sectioning. Precise alignment was carried out to ensure that the objective and the GRIN lens shared the same optical axis.

For two-photon imaging, the fluorescence emission signal collected by the GRIN lens and the objective was directed to the photo-detection unit via dichroic mirror D2 (Semrock, FF705-Di01-25×36). Then the fluorescence was separated by dichroic mirror D3 (Semrock, FF560-Di01-25×36) into a red and a green channel. The band-pass filters F2 (Semrock, FF01-525/50) and F3 (Semrock, FF01-593/46)) were placed before the current photomultiplier tube (PMT) modules (Hamamatsu, H11461-01 and H11461-03) to select a particular wavelength range of detection. The corresponding PMT signal was then fed into a current amplifier (Stanford Research, SR570 and Femto, DLPCA-200) and subsequently into a data acquisition device (National instrument, PCIe-6353) controlled with custom-written software.

For wavefront sensing, D2 was switched to another dichroic mirror (Semrock, Di02-R488-25×36) using a motorized flipper (Thorlabs, MFF101/M) so that the guide star signal of fluorescence emission can transmit through D2 and be descanned by the galvanometer scanning mirrors. Then the fluorescence signal was reflected by the DM and separated from the excitation laser by dichroic mirror D1 (Semrock, FF705-Di01-25×36). The DM was conjugated to the microlens array (SUSS MicroOptics, 18-00197) of the SHWS by the lens pair L9 and L10 (Throlabs, AC254-200-A and AC254-100-A). An ultra-sensitive EMCCD (Andor, iXon3 888) was placed in the focal plane of the microlens array to capture the spot pattern, which provides direct measurement of the wavefront distortion. Overall, the DM, the galvanometer scanning mirrors, the back focal plane of the objective and the microlens array of SHWS are all mutually conjugated via the 4*f* relay system.

### Calibration of the deformable mirror

The DM was calibrated before it was integrated into the microscope system. Following a previously reported procedure^14^, the driving voltage pattern of the DM for the first 65 of Noll’s Zernike modes was measured using a home-built Michaelson interferometer. In this way, the DM can be controlled to take any desired shape via a linear combination of these Zernike modes.

### System AO correction

The aberrations of the microscope system were corrected before any imaging experiment. First, the 10× air objective and the GRIN lens were replaced by another objective with nearly aberration-free performance (Olympus, XLPLN25XWMP2, 25X, 1.05 NA) such that the measured aberrations would mainly come from the imperfections of the optical components or alignments of the system. Then the two-photon fluorescence intensity of a fluorescent dye (rhodamine 6G) solution was used as the feedback to optimize the shape of the DM. Here, a Zernike-mode-based sensorless AO algorithm^15^ was adopted. Briefly, seven to nine different values of each Zernike mode were applied to the DM and the corresponding fluorescence intensity was fitted with a Gaussian function to find the optimal value. This procedure was repeated for the first 21 Zernike modes (tip, tilt and defocus excluded) and the system aberration ***Z***_sys_ could be measured and compensated.

### Calibration of the wavefront sensor

The SHWS was calibrated with respect to the DM in the microscope system. The femtosecond laser was focused by the 25× water objective to excite a fluorescent dye (rhodamine 6G) solution, which created a nonlinear fluorescent guide star. The light of the guide star was shaped by the DM and then sent to the SHWS, which formed the closed-loop AO system. The influence matrix ***M***_sz_ of the DM on the SHWS was calibrated by sequentially applying the first 65 Zernike modes to the DM and recording the corresponding spot displacement in the SHWS. The rows of ***M***_sz_ represent the shift in spot position on the SHWS in response to each Zernike mode and formed the basis on which the *in vivo* wavefront measurement can be decomposed. In this work, we chose the modal wavefront reconstruction algorithm because 1) the aberration is averaged over a small volume and thus is mainly of low order; 2) this algorithm is robust to the noise and missing spots in the *in vivo* wavefront measurement^14^; and 3) each Zernike mode has well defined physically meanings such as astigmatism, coma, and spherical aberration, making the interpretation of GRIN lens aberrations explicit.

### Full AO correction

The full AO correction started with the system aberration corrected and further compensated for the aberrations of the GRIN lens and those induced by biological samples. The reference spot pattern on SHWS ***S***_ref_ = (*x*_1_ ⋯ *x*_*N*_, *y*_1_ ⋯ *y*_*N*_) was recorded using the fluorescence signal of rhodamine at the FOV center of the endomicroscope with aberration corrected by the sensorless algorithm described above. For *in vivo* aberration measurement, the femtosecond laser was scanned over a small FOV (30 μm × 30 μm) and the excited fluorescence signal was descanned and integrated at the SHWS. The spot location of the *in vivo* SHWS measurement ***S***_all_ was estimated using a Gaussian-fit centroid algorithm^16^ and the reliability weight of each spot was determined by its signal-to-background ratio ***W*** = Diag(*w*_1_ ⋯ *w*_*N*_, *w*_1_ ⋯ *w*_*N*_). The spot displacement of the *in vivo* measurement relative to the reference position was calculated as Δ***S*** = ***S***_all_ − ***S***_ref_. Then the additional corrective pattern of the DM could be computed by minimizing the total aberration as follows:

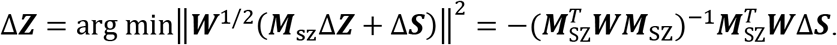

The full AO correction pattern applied to the DM was ***Z***_full_ = ***Z***_sys_ + Δ***Z***. In this paper, all of the plotted wavefront distortions represent the aberrations induced by the GRIN lens and biological samples, i.e. Δ***Z***.

### Animal preparation

C57 (C57BL/6J) and Thy1-GFP (Tg(Thy1-EGFP)MJrs/J) mice were obtained from the Jackson Laboratory. The mice were housed at the Animal and Plant Care Facility of the Hong Kong University of Science and Technology (HKUST). Mice of the same sex were housed four per cage with a 12-h light/dark cycle, and food and water *ad libitum*. All animal procedures were conducted in accordance with the Guidelines of the Animal Care Facility of HKUST and approved by the Animal Ethics Committee at HKUST.

#### Virus injection

The AAV9-CaMKII-GCaMP6f virus was obtained from the Penn Vector Core at the University of Pennsylvania. Virus was diluted 1:10 in PBS and delivered as a bolus (0.5 μL) at 50–100 nL min^−1^ via a Hamilton syringe into the hippocampal CA1 region (anteroposterior, −2.00 mm; mediolateral, ±1.50 mm; dorsoventral, −1.4 mm; relative to the bregma) in 3–4-month-old C57 mice. After injection, the needle was kept in place for 10 min and then retracted from the brain. The mice were then returned to their home cages to recover for 3-4 weeks.

#### GRIN lens implantation

Dexamethasone (0.2 μg/mg) solution was subcutaneously administered one hour before the procedure to prevent brain swelling and reduce inflammatory responses. The mouse was then secured on a stereotaxic instrument and anesthetized with isoflurane (1.5-2% in oxygen). After the skull was exposed and cleaned with ethanol (70%), three stainless steel screws were implanted in the skull to form a triangle with the center located over the hippocampus CA1 region for skull stabilization. Then a 1.6-mm-diameter craniotomy was performed at stereotactic coordinates (2 mm, 1.5 mm) posterior and lateral to the bregma point. Next, the cylindrical column of the exposed cortex was slowly aspirated and removed using a 27-gauge blunt needle until the white matter appeared. The GRIN lens was then positioned over the cranial window and gently inserted into the brain. After a thin layer of adhesive luting cement applied to the skull surrounding the window had dried and hardened, a small amount of dental cement was used to cover the exposed skull surface. A custom-designed rectangular head plate with a round hole of clearance for GRIN lens at the center was then permanently glued to the exposed skull. It would be rigidly mounted the mouse head on a holding device with angle adjusters (NARISHIGE, MAG-2) during *in vivo* imaging experiments. After the surgery, the mouse was allowed to recover from the anesthesia on a heating blanket before returning to its home cage.

### Alignment of the GRIN lens

For *in vitro* and *in vivo* experiments, the sample was mounted on the rotational stage MAG-2 and further fixed to a three-axis translation stage, which enabled precise adjustment of the angle and position of the GRIN lens. A visible laser diode was introduced to the endomicroscopy system to assist the alignment. The laser beam was first aligned with the optical axis of the objective. Then the objective was removed and the tilt/tip of the GRIN lens was adjusted so that the laser was normally incident on the upper surface of the GRIN lens. This guaranteed that the optical axes of the GRIN lens and the objective were parallel to each other. Next, we looked at the imaging plane of the objective through the eyepiece and translated the GRIN lens until its upper surface came into focus. Then the objective was lifted 100 μm up so that the GRIN lens could operate at its designed working distance. Finally, we switched to the two-photon imaging mode and translated the GRIN lens sideways until the fluorescence image appeared at the center of the FOV.

### Lookup table for the aberrations of the GRIN lens

To evaluate the efficacy of our approach, we first characterized the intrinsic aberrations of the GRIN lens that was embedded in a rhodamine/agarose mixture that contains sparsely distributed and immobilized fluorescent beads (200-nm diameter). Its aberration was measured by the wavefront distortion of rhodamine fluorescence, whereas the PSF was determined by imaging the individual fluorescent bead. To achieve the optimal imaging performance, the GRIN lens was precisely aligned to share the same optical axis with the objective. Considering the cylindrical symmetry of the GRIN lens, the cylindrical coordinates (r, θ, z) were adopted to describe the imaging location in the sample where z=0 corresponds to the designed focal plane of the GRIN lens. The representative on-axis and off-axis aberrations of the GRIN lens at various depths are shown in **Supplementary Figs. 2 and 3**, which are primarily dominated by spherical aberration and astigmatism respectively, consistent with a previous study^5^. As can be seen from **Supplementary Figs. 2 and 3**, despite the aberrations increasing rapidly and the PSF becoming severely distorted when the imaging position deviates from the center point, AO correction can effectively increase the fluorescence signal and recover the diffraction-limited performance at various depths. Because the optical properties of the GRIN lens are determined by the optical design and manufacturing process, we can pre-calibrate the intrinsic aberrations of the GRIN lens and store them in a lookup table. The lookup table was established using an *in vitro* setup. The detailed imaging parameters were listed in **Supplementary Table 1**. In principle, we need to measure all the aberrations at different field locations. Considering that the aberrations of the GRIN lens are cylindrically symmetrical along the optical axis^4,5^, the number of aberration measurements could be greatly reduced. To cover the entire cylindrical FOV with a radius of 150 μm and a depth of 300 μm, we built the lookup table by measuring the aberrations along the *r* and *z* directions at 30 μm intervals, resulting in 6×11 wavefront distortions in the *θ* = 0 subplane (**Supplementary Fig. 6a**).

For *in vivo* imaging, the photon budget is limited and the two-photon fluorescence intensity was practically too weak for wavefront measurement, especially in the off-axis region of large aberration. To solve this problem, we precompensated for the aberration of the GRIN lens in the imaging region (*r*_1_, *θ*_1_, *z*_1_) using the lookup table (**Supplementary Fig. 6b**). Firstly, the aberration of the GRIN lens at location (*r*_1_, 0, *z*_1_) was estimated from the linear interpolation of the aberrations nearby stored in the lookup table. Secondly, the aberration at the center was subtracted from that at (*r*_1_, 0, *z*_1_). Lastly, the resulting aberration was rotated by the angle *θ*_1_ and added to the aberration at the center, which was the estimated wavefront correction for the ROI (*r*_1_, *θ*_1_, *z*_1_). As shown in **Fig. 3**, because of the intrinsic aberrations of the GRIN lens were largely corrected by using the lookup table, the fluorescence intensity and imaging resolution were improved tremendously. Then we were able to perform *in vivo* AO correction based on direct wavefront sensing to further eliminate the residual aberrations caused by the alignment error of the GRIN lens as well as biological tissues.

### *In vivo* imaging

The imaging experiment was performed three weeks after the implantation of the GRIN lens. The mouse was first briefly anesthetized with isoflurane (1-2% in oxygen) and mounted on the head-holding device. The GRIN lens was precisely aligned to share the same optical axis with the objective lens. The femtosecond laser tuned at 920 nm was used to excite GFP and GCaMP6s. For wavefront sensing, the aberration of the GRIN lens was first precompensated using the lookup table and the residual aberration was measured and corrected using the two-photon excited guide star. The laser power was 30~50 mW through the objective and the exposure time was less than 2 s. Detailed imaging parameters including excitation power, pixel rate, guide star integration time can be found in **Supplementary Table 2-3**. For the morphological imaging, the animals were anesthetized with isoflurane (1% in oxygen) with the head secured on a head-holding stage (NARISHIGE, MAG-2). For calcium imaging, the mice were fully awake or lightly anesthetized with isoflurane and placed on a head-fixation behavioral platform that was based on a rotary treadmill.

### Laser-mediated neuronal injury

The field-dependent GRIN lens aberration was first measured and corrected as previously described to enable precise and efficient laser-mediated microsurgery. For laser cutting of the dendritic branch, the 920 nm laser with 160-mW power through the objective was line scanned (1s/line) across the branch repeatedly for 20 second. For micro-lesion of dendritic shaft, the laser with the same optical power was focused on the shaft center for 5 second. The parameters of optical power and duration for laser microsurgery have been optimized to induce reproducible neuronal injuries.

### Induction of epileptic seizure

After the baseline image was obtained, the mice were allowed to recovered from the isoflurane anesthesia. Then single high dose (20mg/kg) of kainic acid (Simga-Aldrich) dissolved in Dulbecco’s Phosphate-Buffered Saline (DPBS) was intraperitoneally (i.p.) injected to fully awake mice and the mice status epilepticus was quantified by the Racine stages17. After the mice progressed to stage 3 or above, they were allowed to stayed in the cage for 30 minutes before the seizure was terminated by isoflurane anesthesia. For control experiment, the mice received DPBS injection instead of kainic acid.

### Multiplane calcium imaging

An electrically tunable lens (Optotune, EL-16-40-TC-VIS-5D-C) was placed close to the rear stop of the 10X objective lens for quick tuning of the focal plane. We first picked three ROIs and measured the wavefront distortion using the method described above. The three corrective patterns of the DM were stored locally for further use. Next, we conducted near-simultaneously calcium imaging of the three ROIs at a volume rate of 5 Hz. The ETL and DM were synchronized with the scanner as shown in **Supplementary Fig. 9**. The driving voltage of the ETL was low-pass filtered to avoid high-order oscillation and improve the temporal response. The pre-measured corrective patterns were updated on the DM accordingly to compensate for the field-dependent aberrations.

### Image analysis

The images were processed in Matlab or ImageJ^18^. For image deconvolution, the Richardson-Lucy algorithm of the DeconvolutionLab2 plugin in ImageJ was adopted. The PSF used for deconvolution was experimentally measured at different imaging positions. To make the PSF measurement less laborious, we took advantage of the cylindrical symmetry of the GRIN lens and only measured the PSF in the *θ* = 0 subplane. The PSF in other regions could be estimated using the rotation-based procedure. For the mosaic images, multi-tile subimages were stitched together using either MosaicJ^19^ or Grid/Collection Stitching^20^ plugin in ImageJ. To cover the entire FOV of 300 μm in diameter, 81 subimages at 30 μm intervals were acquired.

## Supplementary information

**Supplementary Figure 1.**
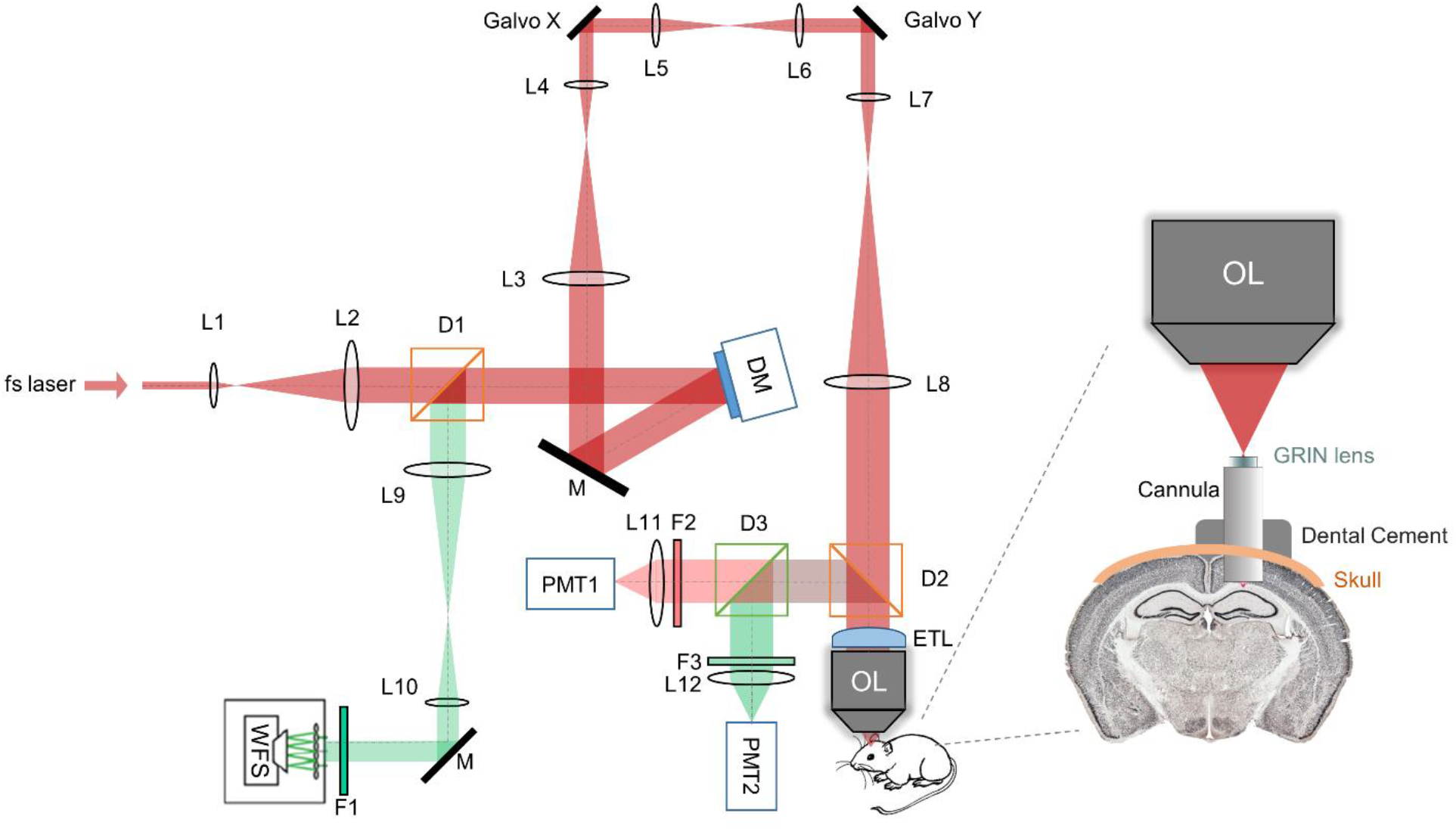
Schematic diagram of our AO two-photon endomicroscope system for *in vivo* deep brain imaging. L1-L12: lenses; OL: objective lens; D1-D3: dichroic mirrors; F1-F3: filters; M: mirrors; DM: deformable mirror; WFS: wavefront sensor; PMT1-2: photomultiplier tubes; ETL: electrically tunable lens.

**Supplementary Figure 2.**
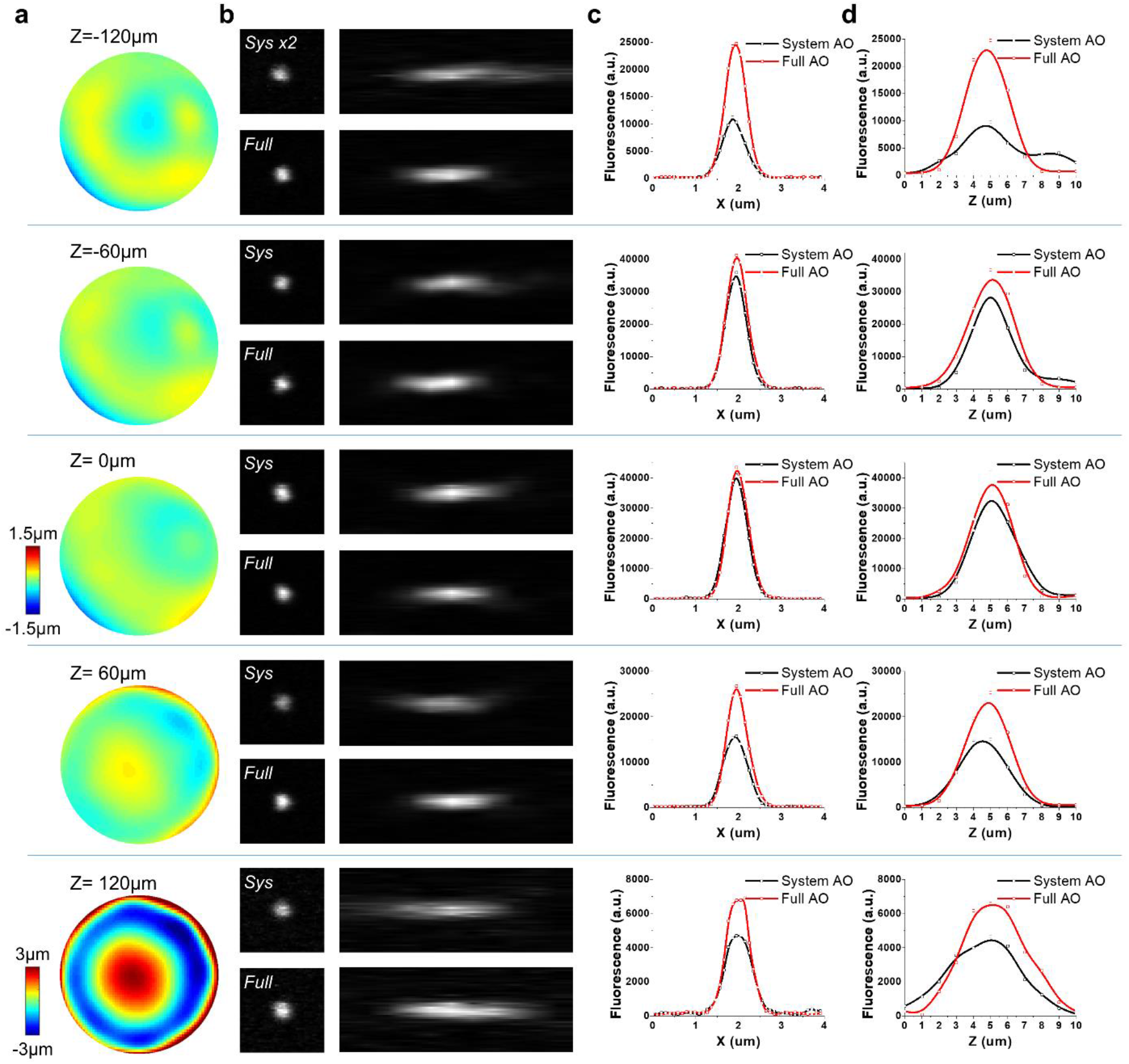
Characterization of the on-axis aberrations of the GRIN lens. Column (a): The wavefront distortion at different depths along the optical axis. Column (b): Lateral and axial PSF measured with fluorescent beads that were 0.2 μm in diameter at the corresponding depths. Column (c): The lateral intensity profile before and after AO correction at the corresponding depths. Column (d): The axial intensity profile before and after AO correction at the corresponding depths.

**Supplementary Figure 3.**
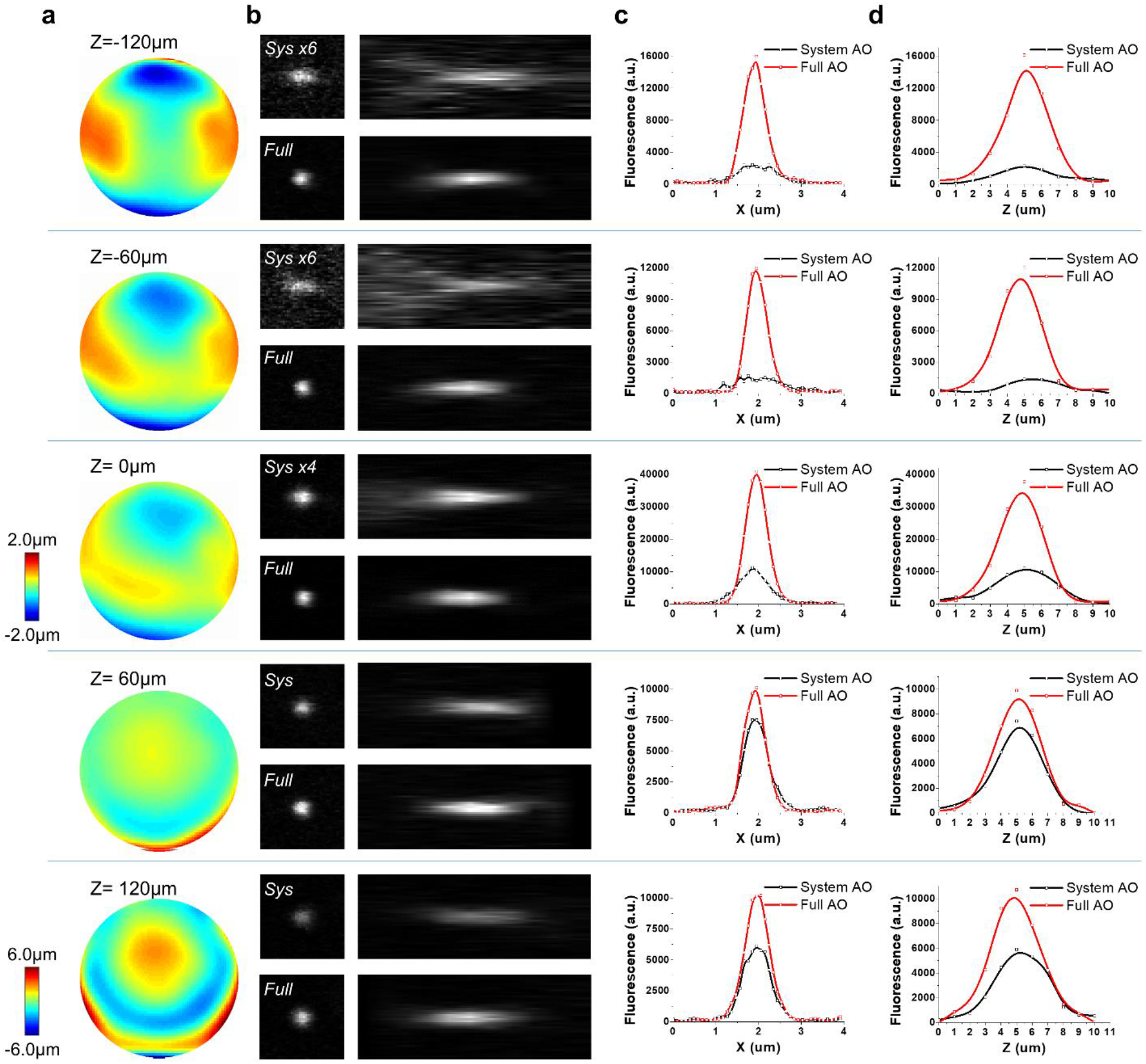
Characterization of the off-axis (r=60 μm) aberrations of the GRIN lens. Column (a): **(a)** The wavefront distortion at a distance of 60 μm from the field center at different depths along the optical axis. Column (b): Lateral and axial PSF measured with fluorescent beads that were 0.2 μm in diameter at the corresponding depths. Column (c): The lateral intensity profile before and after AO correction at the corresponding depths. Column (d): The axial intensity profile before and after AO correction at the corresponding depths.

**Supplementary Figure 4.**
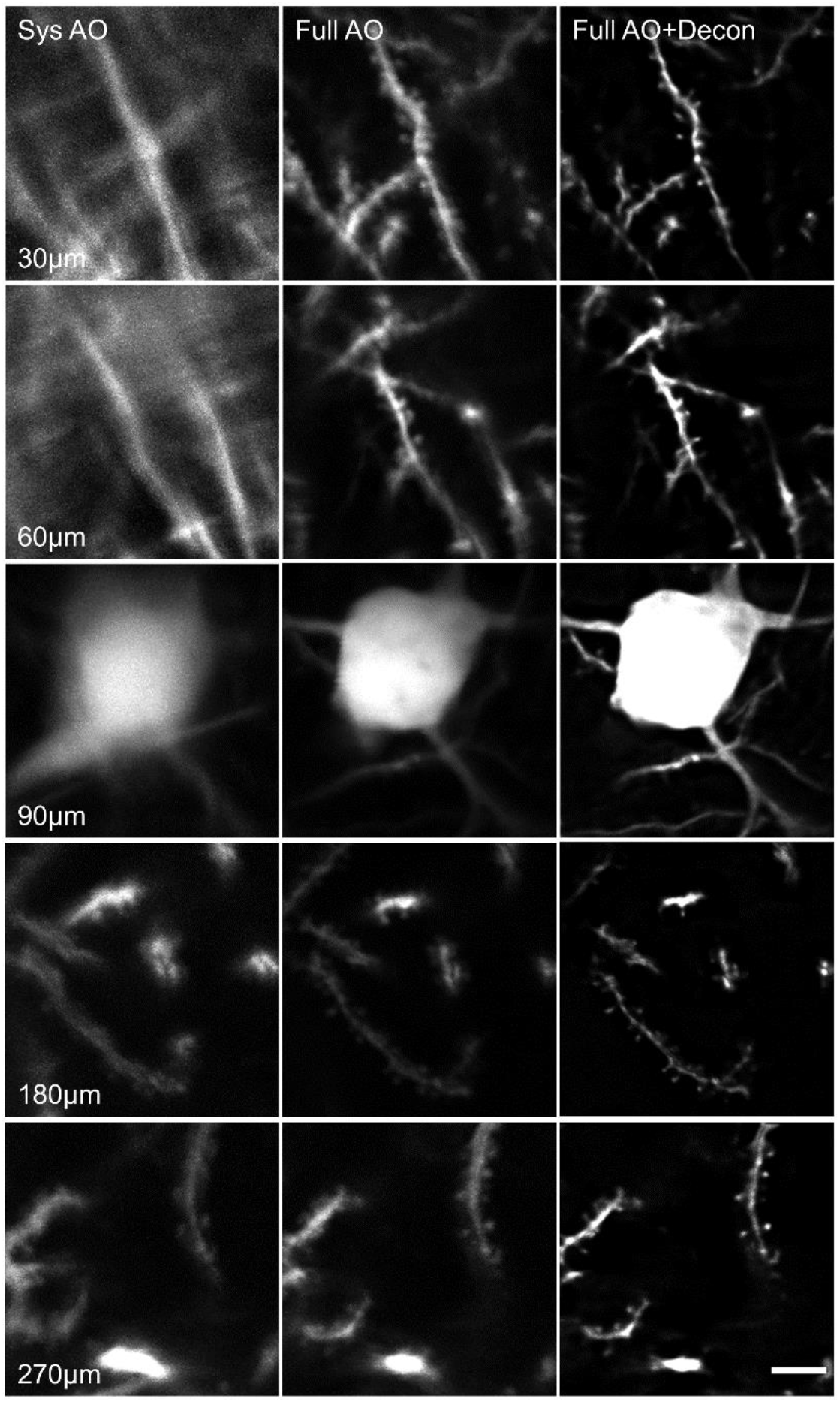
Comparison of system AO, full AO and full AO with deconvolution at different imaging depths. The results demonstrate that AO can significantly improve the imaging resolution at all imaging depths. Scale bar: 5 μm.

**Supplementary Figure 5.**
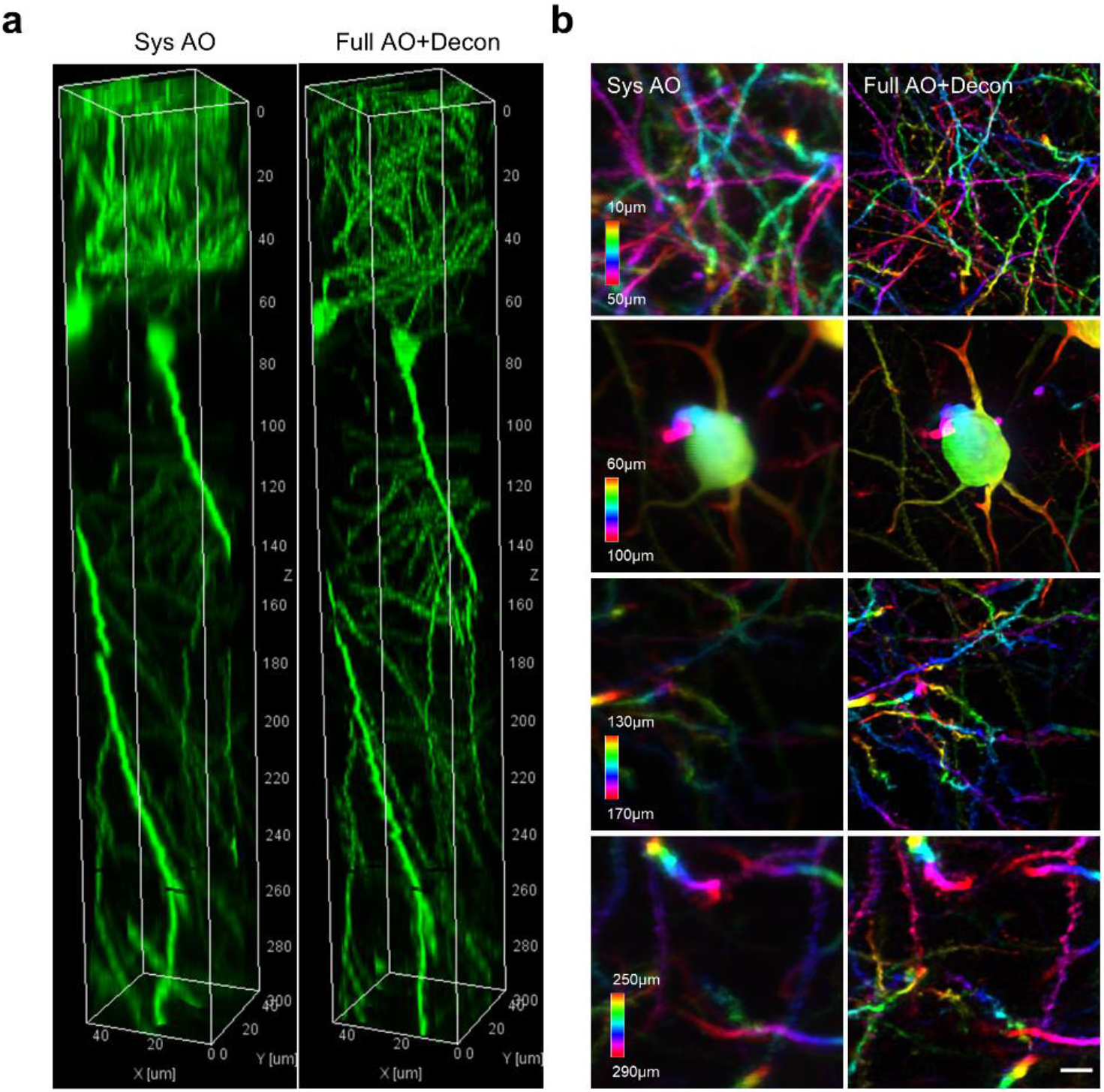
Three-dimensional imaging of the column located at the center (r=0). **(a)** In vivo imaging of GFP-labeled CA1 neurons in a 50×50×300 μm^3^ volume with system (left) and full (right) AO correction. **(b)** MIPs of four subvolumes with system (left) and full (right) AO correction where the structures are color-coded by depth. Scale bar: 5 μm.

**Supplementary Figure 6.**
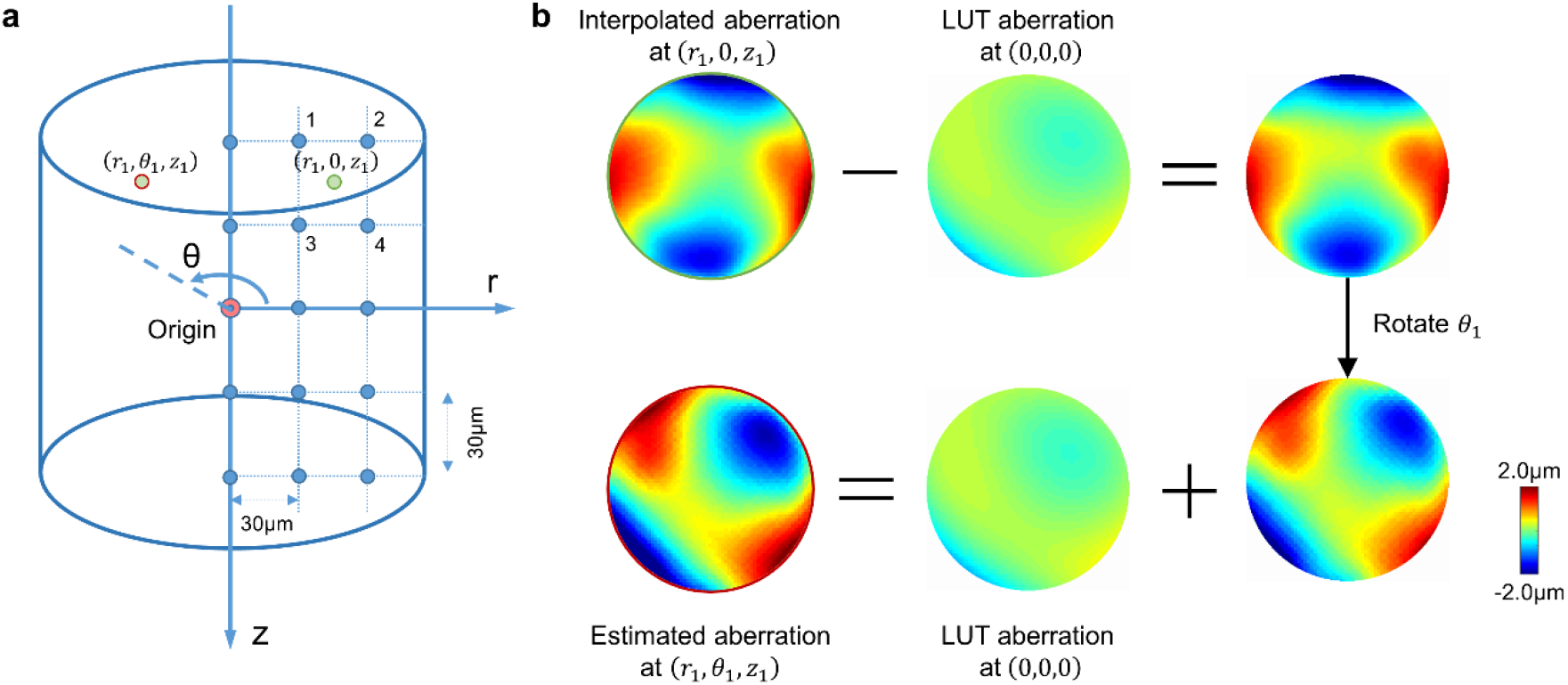
Lookup table for the aberrations of the GRIN lens. **(a)** Calibration of the lookup table. Considering the cylindrical symmetry of the GRIN lens, the cylindrical coordinate system was used to describe the imaging location. The origin is defined as the center located at the designed working distance of the GRIN lens. The entire imaging FOV had a radius of 150 μm and a depth of 300 μm. We measured the intrinsic aberration of the GRIN lens in the *θ*=0 subplane at 30 μm intervals, shown as blue dots in the figure. **(b)** Estimation of GRIN lens-induced aberration using the lookup table. To find the aberration at location (*r*_1_, *θ*_1_, *z*_1_) (red dot in (a)), we first estimate the aberration at the rotational symmetrical point (*r*_1_, 0, *z*_1_) using linear interpolation of the aberrations nearby (the blue dots labeled 1-4). Then the aberration at the origin is subtracted from the interpolated wavefront distortion, and the resulting wavefront is rotated by the angle *θ*_1_ and added back to the aberration at the origin.

**Supplementary Figure 7.**
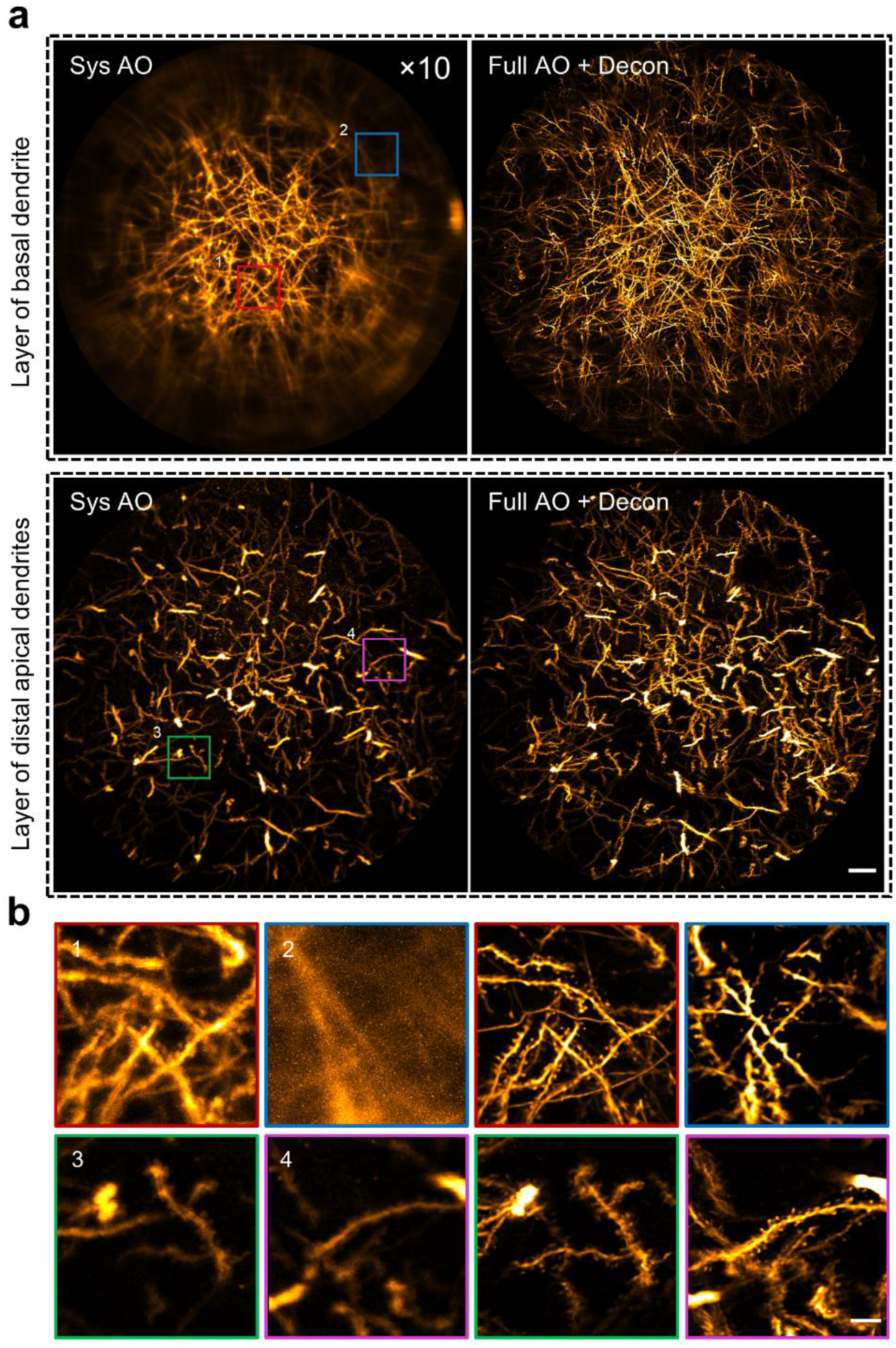
AO improves resolution over a large FOV. **(a)** MIPs of the basal dendrite layer and the distal apical dendrite layer with system (left) and full (right) AO correction. The entire FOV is 300 μm in diameter. Scale bar: 20 μm. **(b)** Left: four magnified views of the sub-regions indicated by the numbered boxes in (a). Depth range of projection for the layer of basal dendrites: 20-50 μm; and for layer of distal apical dendrites: 190-220 μm. Scale bar: 5 μm.

**Supplementary Figure 8.**
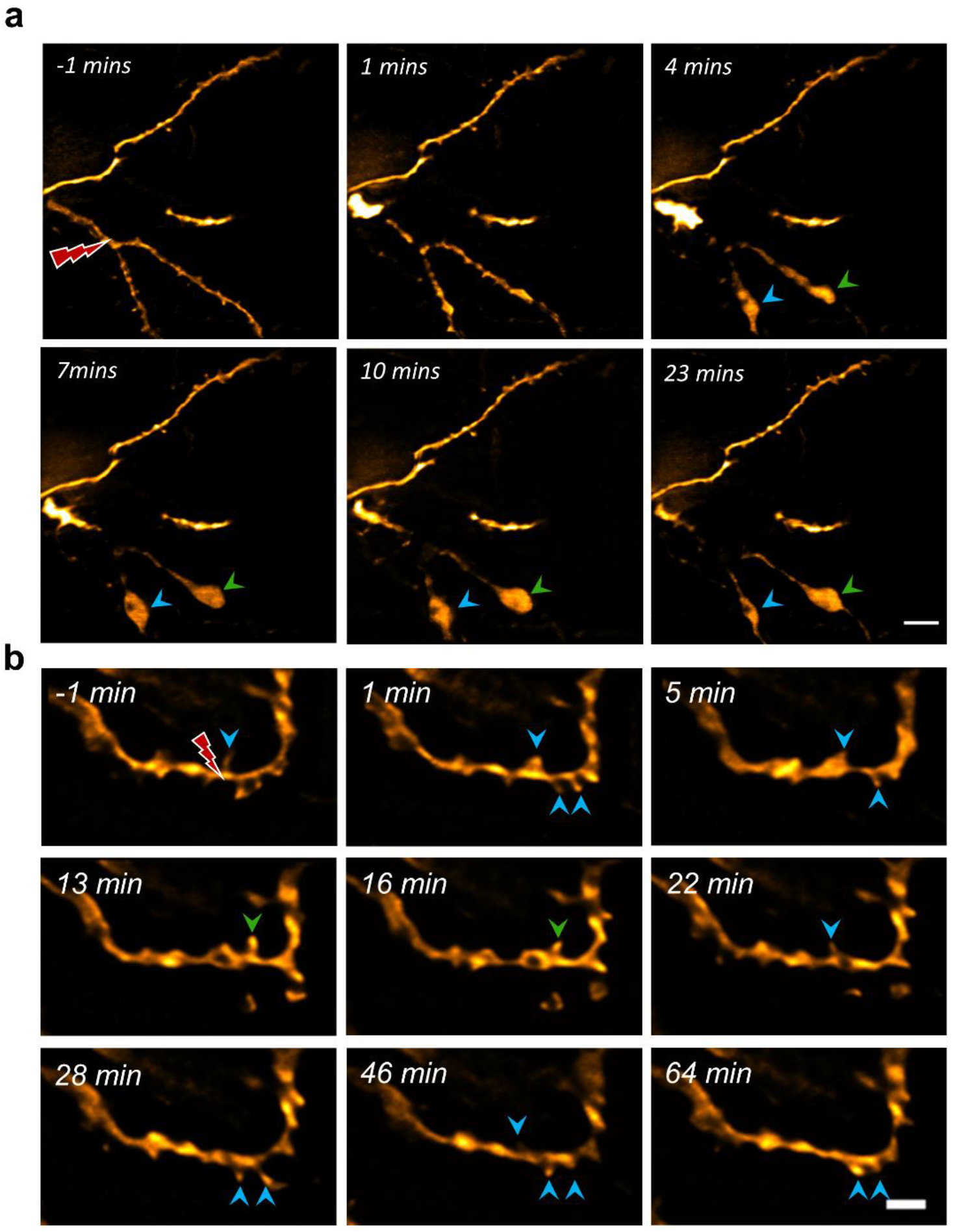
Time-lapse imaging of dendritic dynamics after laser injury. (a) Laser cutting of dendritic branches leads to rapid and prolonged degeneration that resembles Wallerian axonal degeneration. The bead-like formation at the distal end of the dendrites were indicated by the colored arrowheads. Scale bar: 5 μm (b) Laser-mediated micro-lesion of dendritic shaft causes recurrent spinogenesis near the site of injury. The blue and green arrowhead indicate recurrent spine and newly-appeared spine, respectively. Scale bar: 2 μm.

**Supplementary Figure 9.**
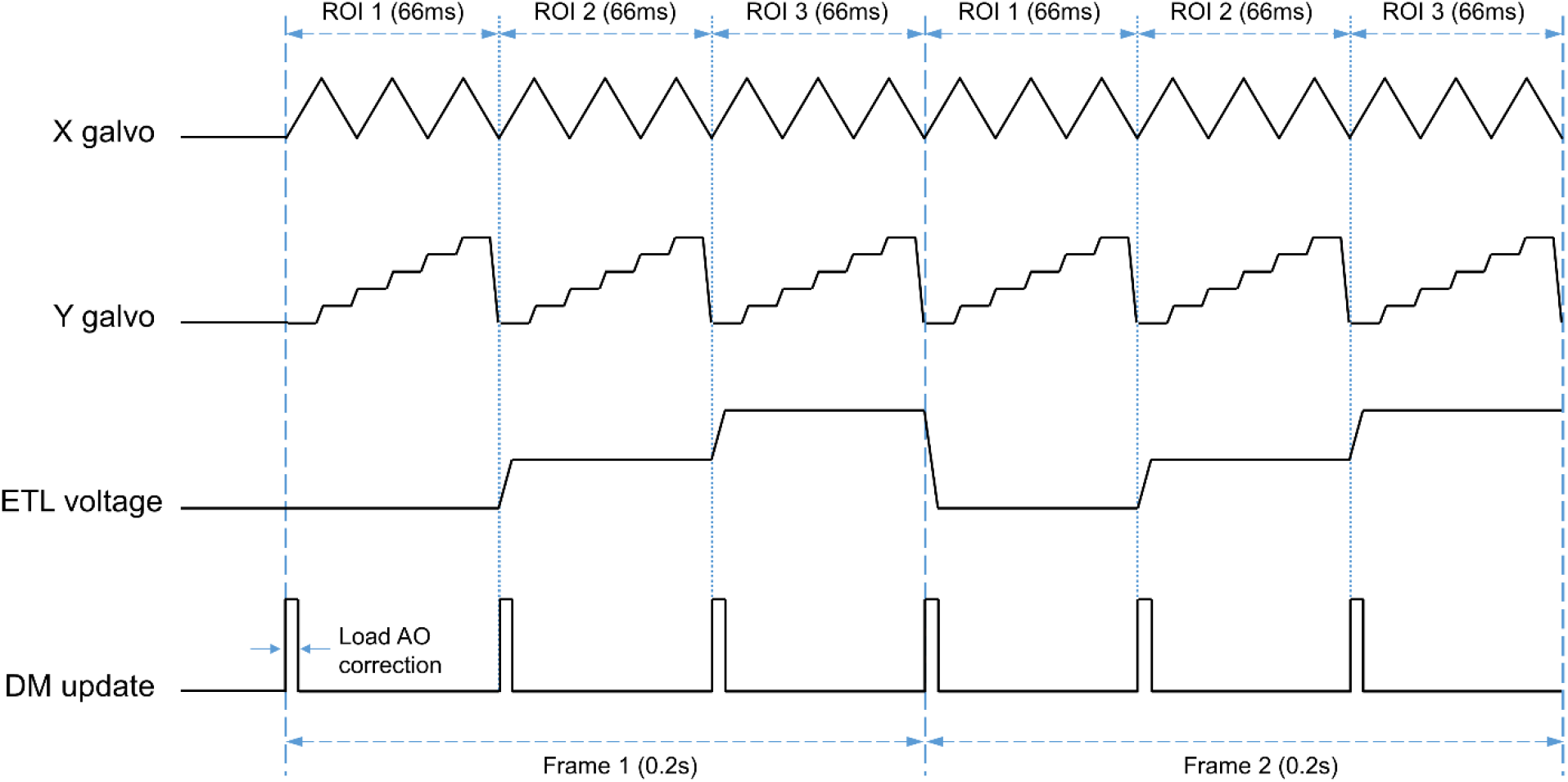
Control signals of x, y scanners, the ETL and the DM for random-access multiplane imaging.

**Supplementary Figure 10.**
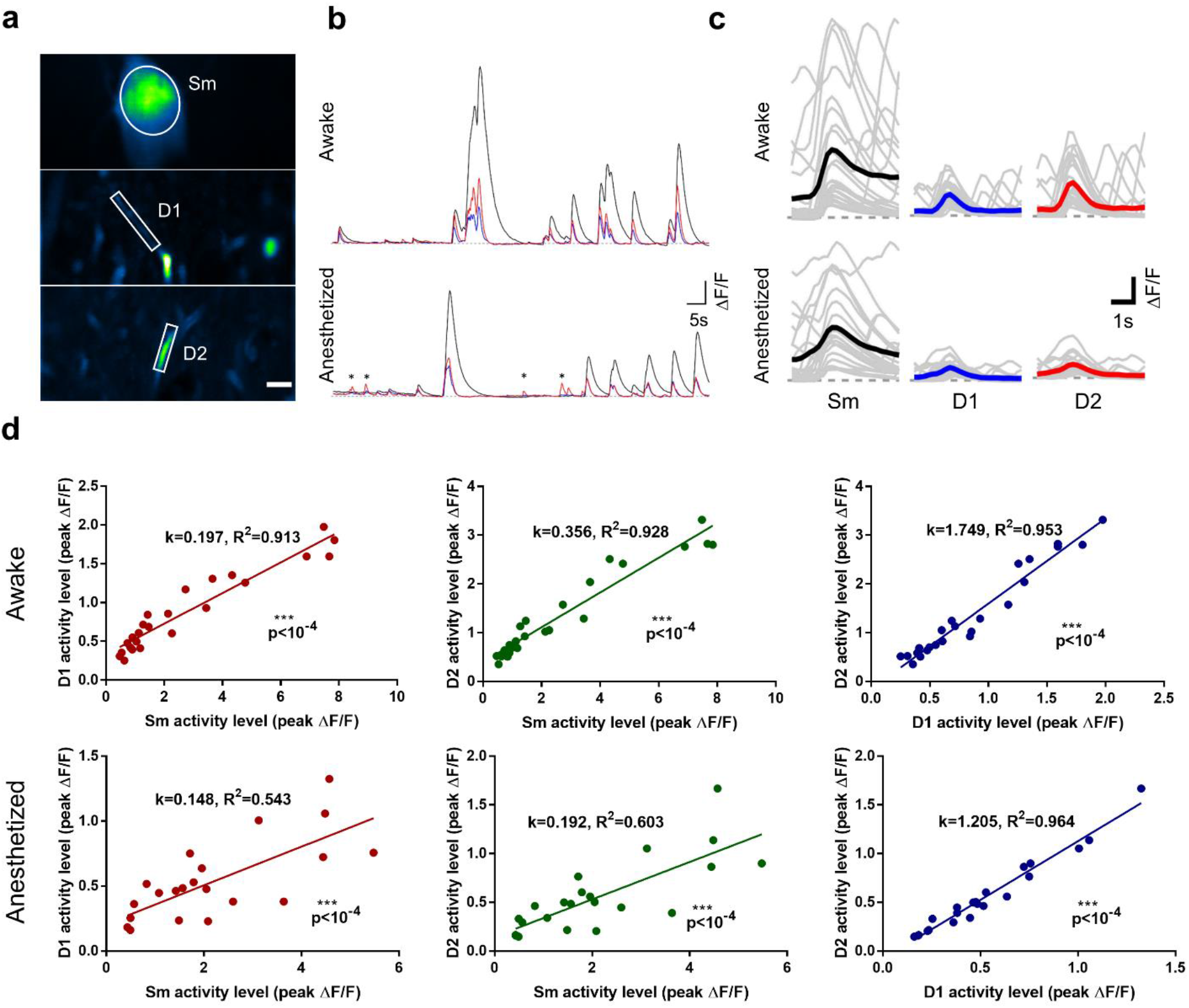
Correlated somato-dendritic spontaneous activity of hippocampal CA pyramidal neuron is brain-state dependent. (a) *In vivo* multiplane calcium imaging of somato (Sm) and dendritic (D1, D2) activity of single neuron in mice hippocampus. Images are shown as STD projection of the 600 frames. Scale bar: 5μm. (b) Calcium transients (ΔF/F) of soma and dendrites shown in (a) under mice wakefulness (upper) and isoflurane-induced anesthesia (lower). Black: Sm, Blue: D1, Red: D2. The asterisks indicate the unpaired dendritic transients appeared in isoflurane anesthesia. (c) Firing events of soma (Sm) and dendrites (D1, D2) at different brain states. Grey and colored curves represent the individual and average event, respectively. (d) Relationship between activity strength of soma and dendrites under mice wakefulness and anesthesia.

**Supplementary Table 1.**
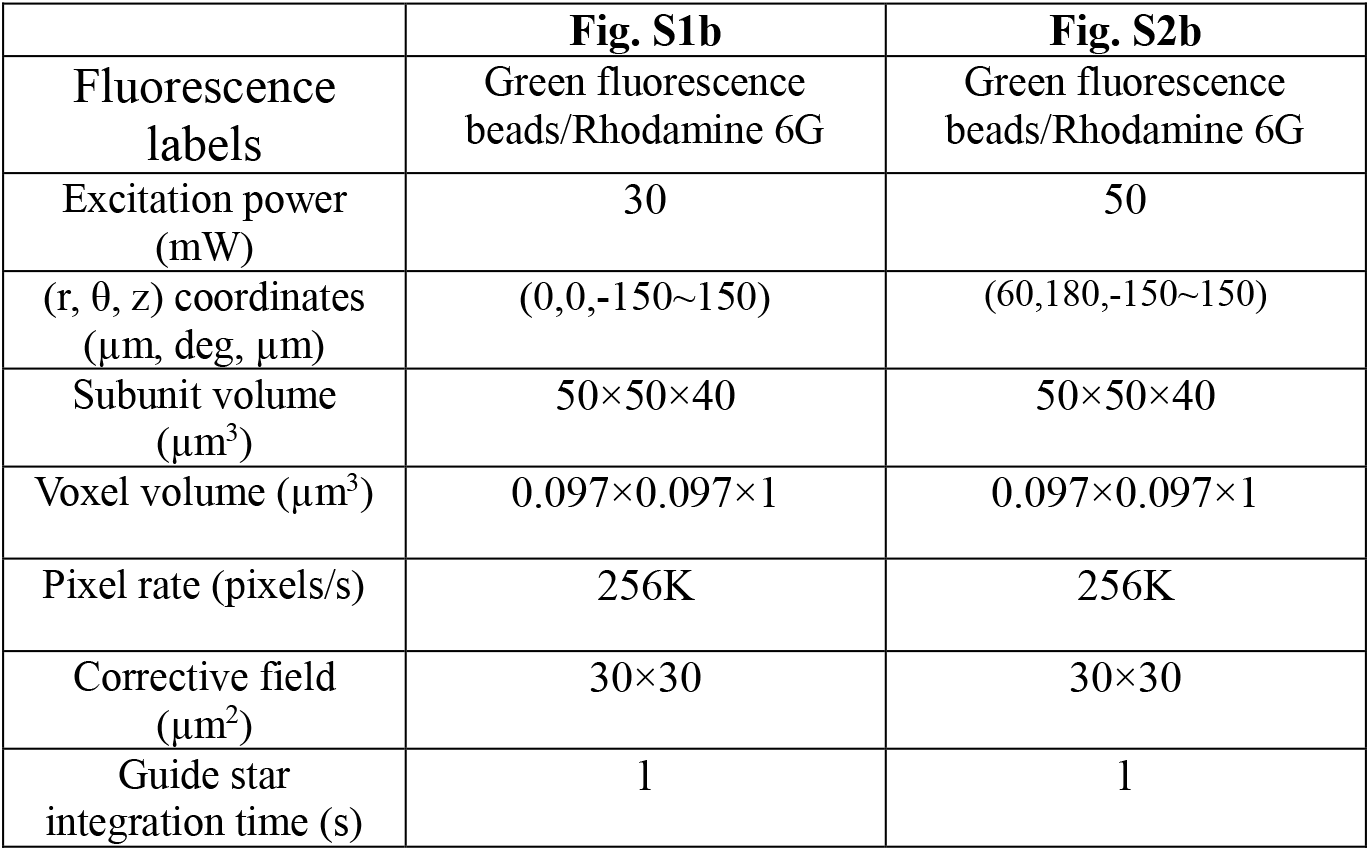
Wavefront sensing and in vitro imaging parameters.

|  | **Fig. S1b** | **Fig. S2b** |
| --- | --- | --- |
| Fluorescence labels | Green fluorescence beads/Rhodamine 6G | Green fluorescence beads/Rhodamine 6G |
| Excitation power (mW) | 30 | 50 |
| (r, $\theta$, z) coordinates ($\mu\text{m}$, deg, $\mu\text{m}$) | (0,0,-150~150) | (60,180,-150~150) |
| Subunit volume ($\mu\text{m}^3$) | 50×50×40 | 50×50×40 |
| Voxel volume ($\mu\text{m}^3$) | 0.097×0.097×1 | 0.097×0.097×1 |
| Pixel rate (pixels/s) | 256K | 256K |
| Corrective field ($\mu\text{m}^2$) | 30×30 | 30×30 |
| Guide star integration time (s) | 1 | 1 |

**Supplementary Table 2.** Wavefront sensing and *in vivo* morphological imaging parameters.

|  | **Fig. 1a** | **Fig. S4a** | **Fig. S5a** | **Fig. S8a** |
| --- | --- | --- | --- | --- |
| Fluorescence labels | GFP | GFP | GFP | GFP |
| Excitation power (mW) | 30 | 50 | 30 | 30 |
| (r, $\theta$, z) coordinates ($\mu\text{m}$, deg, $\mu\text{m}$) | (60, -162, -150~150) | (96,106, -150~150) | (0,0,-150~150) | (0,0,-95--55) |
| Subunit volume ($\mu\text{m}^3$) | 50×50×40 | 50×50×40 | 50×50×40 | 50×50×40 |
| Voxel volume ($\mu\text{m}^3$) | 0.097×0.097×1 | 0.097×0.097×1 | 0.097×0.097×1 | 0.097×0.097×1 |
| Pixel rate (pixels/s) | 256K | 256K | 256K | 256K |
| Corrective field ($\mu\text{m}^2$) | 30×30 | 30×30 | 30×30 | 30×30 |
| Guide star integration time (s) | 1~2 | 1~3 | 1~2 | 1 |

**Supplementary Table 2 (continued) | Wavefront sensing and *in vivo* morphological imaging parameters.**
|  | **Fig. 2a (upper)** | **Fig. 2a (lower)** | **Fig. S7a (upper)** | **Fig. S7a (lower)** |
| --- | --- | --- | --- | --- |
| Fluorescence labels | GFP | GFP | GFP | GFP |
| Excitation power (mW) | 30 | 30 | 30 | 50 |
| (r, $\theta$, z) coordinates ($\mu\text{m}$, deg, $\mu\text{m}$) | (0~150, 0~360, -90~-50) | (0~150, 0~360, -20~20) | (0~150, 0~360, -140~-100) | (0~150, 0~360, 30~70) |
| Subunit volume ($\mu\text{m}^3$) | 50×50×40 | 50×50×40 | 50×50×40 | 50×50×40 |
| Voxel volume ($\mu\text{m}^3$) | 0.097×0.097×1 | 0.097×0.097×1 | 0.097×0.097×1 | 0.097×0.097×1 |
| Pixel rate (pixels/s) | 984K | 128K | 128K | 256K |
| Corrective field ($\mu\text{m}^2$) | 30×30 | 30×30 | 30×30 | 30×30 |
| Guide star integration time (s) | 1 | 1 | 1 | 1 |

**Supplementary Table 3.** Wavefront sensing and *in vivo* calcium imaging parameters.

|  | **Fig. 4b** | **Fig. 5b** | **Fig. S4d** | **Fig. S10a** |
| --- | --- | --- | --- | --- |
| Fluorescence labels | GCaMP6s | GCaMP6s | GCaMP6s | GCaMP6s |
| Excitation power (mW) | 30 | 30 | 30 | 30 |
| (r, $\theta$, z) coordinates ($\mu\text{m}$, deg, $\mu\text{m}$) | P1: (84, -71, -99)<br>P2: (80, -55, -42)<br>P3: (79, 78, -78) | P1: (156, -88, -108)<br>P2: (79, 78, -83)<br>P3: (61, 81, -57) | P1: (64, -1.7, -119)<br>P2: (93, -83, -119)<br>P3: (86, 82, -109) | P1: (156, -88, -108)<br>P2: (80, 78, -80)<br>P3: (64, 81, -53) |
| FOV of each plane ($\mu\text{m}^2$) | 50×26 | 50×26 | 50×26 | 50×26 |
| Pixel size ($\mu\text{m}^2$) | 0.195×0.195 | 0.195×0.195 | 0.195×0.195 | 0.195×0.195 |
| Pixel rate (pixels/s) | 507K | 507K | 507K | 507K |
| Corrective Field ($\mu\text{m}^2$) | 30×30 | 30×30 | 30×30 | 30×30 |
| Guide star integration time (s) | 1 | 1 | 1 | 1 |

**Supplementary Video 1 | AO enables multiplane Ca^2+^ imaging of pyramidal neuron at synaptic resolution in awake behaving mice.** Three planes (P1-3) were sequentially captured at 5Hz for calcium imaging of neuronal somata, dendrites and spines, respectively, in the hippocampus CA1 during mice wakefulness.

**Supplementary Video 2 | AO enables accurate recording of calcium transients by eliminating the cross talk of neighboring neurons.** Three ROIs in the stratum pyramidale of hippocampus CA1 were sequentially captured at 5Hz. With AO full correction, the neighboring neurons can be distinguished from each other without cross talk of the fluorescence signal.

**Supplementary Video 3 | Simultaneous Ca^2+^ imaging of somato and dendritic activity of single pyramidal neuron in awake mice.** The Soma and two pyramidal dendrites of a pyramidal neuron in hippocampus CA1 were selected for near simultaneous Ca^2+^ imaging. It shows that somato and dendritic activity are highly correlated.

